# A quantitative PCR based environmental DNA assay for detecting Atlantic salmon (*Salmo salar* L.)

**DOI:** 10.1101/226829

**Authors:** Siobhán Atkinson, Jeannette E.L. Carlsson, Bernard Ball, Damian Egan, Mary Kelly-Quinn, Ken Whelan, Jens Carlsson

## Abstract

1. The Atlantic salmon (*Salmo salar* L.) has worldwide ecological, cultural and economic importance. The species has undergone extensive decline across its native range, yet concerns have been raised about its invasive potential in the Pacific. Knowledge on the distribution of this species is vital for addressing conservation goals.
2. This study presents an eDNA assay to detect *S. salar* in water samples, using quantitative PCR (qPCR) technology. Species-specific primers and a minor groove binding (MGB) probe were designed for the assay, based on the mitochondrial cytochrome oxidase I (COI) gene.
3. The results of this study indicate that eDNA is a highly sensitive tool for detecting *S. salar in situ*, and could potentially provide an alternative, non-invasive method for determining the distribution of this species.

## 1. Introduction

The Atlantic salmon, (*Salmo salar* L.), is of ecological, cultural and economic importance. As a result, this species has been the subject of intense exploitation ranging from commercial fisheries, recreational fishing and intensive aquaculture (Morton, Ariza, Halliday, & Pita, 2016; Piccolo & Orlikowska, 2012). Although *S. salar* is protected under Annex II and Annex V of the EU Habitats Directive, and efforts to reduce fishing pressure and restore freshwater habitats have been implemented, this once abundant species has continued to decline (Chaput, 2012; Friedland et al., 2009). Numerous factors including recruitment failure at sea (Chaput, 2012; Friedland et al., 2009), obstacles to migration in freshwater (Thorstad, Økland, Aarestrup, & Heggberget, 2008) and pollution from agricultural, industrial and urban sources (Hendry, Cragg-Hine, O’Grady, Sambrook, & Stephen, 2003) have contributed to the deterioration of *S. salar* populations. Furthermore, the species is used for intensive aquaculture outside its native range. Large escapes of *S. salar* happen with regularity in these areas, causing concerns about the species’ invasive potential (Fisher, Volpe, & Fisher, 2014; Piccolo & Orlikowska, 2012). To adequately address these issues, and to achieve the conservation objectives of the species, it is vital to have knowledge on its distribution. At present, *S. salar* monitoring involves electrofishing surveys, the placement of fish counters or traps, rod catch data provided by anglers and redd counts (The Standing Scienctific Commitee on Salmon, 2016). These surveys can be expensive, labour intensive and also potentially harmful to the fish (Snyder, 2004). Clearly, there is a need for an effective, efficient and non-invasive sampling method to monitor the species. To this end, environmental DNA (eDNA) analysis may provide an alternative sampling strategy for monitoring the distribution of *S. salar* for management and conservation purposes. Environmental DNA is the collective term for DNA present freely in the environment which has been shed by organisms (in the form of mucus, faeces, gametes or blood, for example), and can be extracted (Taberlet, Coissac, Hajibabaei, & Rieseberg, 2012; Thomsen & Willerslev, 2015). It has been shown to be an effective method for detecting species in freshwater (Clusa, Ardura, Fernández, Roca, & García-Vázquez, 2017; Gustavson et al., 2015), marine (Gargan et al., 2017) and terrestrial (Willerslev, 2003) environments.

Recent studies have developed and deployed specific primers for the detection of *S. salar* in eDNA water samples. A study by Clusa, Ardura, Fernández, Roca, & García-Vázquez (2017), for example, developed *S. salar*–specific primers using the 16S ribosomal DNA (rDNA) region. These authors successfully identified *S. salar* in their eDNA samples using PCR-RFLP (Polymerase chain reaction-restriction fragment length polymorphism). Alternatively, Dalvin, Glover, Sørvik, Seliussen and Taggart (2010) utilised the mitochondrial DNA (mtDNA) cytochrome c oxidase (COI) gene for their primer development, followed by traditional PCR analysis. While the COI primers in this study were successful in amplifying DNA from tissue samples (both fresh and degraded) the authors were unable to detect *S. salar* DNA in their eDNA samples (Dalvin et al., 2010). The assay presented here provides an improvement on these studies. As well as developing species-specific primers with the COI gene, the present assay incorporates an additional species-specific minor groove binding (MGB) probe which allows the eDNA sample to be analysed in quantitative PCR (qPCR). Furthermore, the MGB probe allows for additional sensitivity and specificity of the assay, as three sequences as opposed to two are checked against the target template DNA (Herder et al., 2014).

The aim of this study was to develop an MGB based qPCR assay to detect the presence of *S. salar*. As observed in other studies (Laramie, Pilliod, & Goldberg, 2015) this approach may also allow for the detection of *S. salar* populations in locations where they have not been recorded with traditional methods.

## 2. Methods

### 2.1 eDNA qPCR assay development

Primer Express 3.0 (Applied Biosystems-Roche, Branchburg, NJ) was used to design the species-specific primers (forward primer: 5’-CGC CCT AAG TCT CTT GAT TCG A-3’, and reverse primer 5’-CGT TAT AAA TTT GGT CAT CTC CCA GA-3’) and 5’ NED labelled TaqMan® minor groove binding probe (5’-AGA ACT CAG CCA GCC TG-3’) for *S. salar*, which targeted the mtDNA COI region. The total amplicon size, including primers, was 74 base pairs. Probe and primer sequences were matched against the National Centre for Biotechnology Information (NCBI - http://www.ncbi.nlm.nih.gov/) nucleotide database with BLASTn (Basic Local Alignment Search Tool) to verify the species specificity for the *in silico S. salar* assay. The *S. salar* assay was tested *in vitro* with both closely related and other fish species (marine and freshwater) including brown trout (*S. trutta*), sea lamprey (*Petromyzon marinus* L.), pink salmon (*Oncorhynchus gorbuscha* Walbaum) and herring (*Clupea harengus* L.) to ensure the assay did not amplify other fish species. The qPCR assay was optimized using tissue extracted from *S. salar*.

### 2.2. Study area and field validation of *S. salar*

Three rivers located in the south of Ireland were selected for field validation of the eDNA assay: the Dalligan, Dinin and Burren rivers (Table 1, Figure 1). Each of these rivers contains an obstacle or barrier, which has the potential to prevent or delay the migration of *S. salar*. Electrofishing was carried out by Inland Fisheries Ireland upstream and downstream of each obstacle in July 2017 to verify the presence or absence of *S. salar* at each site. Environmental DNA samples were collected on the same day that the electrofishing was carried out.

**Table 1.** The different combinations of *S. salar* and *S. trutta* presence/absence downstream and upstream of the river obstacles listed. The occurrence of each species was confirmed by electrofishing.

| River | Obstacle Type | *S. salar* Downstream | *S. salar* Upstream | *S. trutta* Downstream | *S. trutta* Upstream |
| --- | --- | --- | --- | --- | --- |
| Burren | Weir | Yes | Yes | Yes | Yes |
| Dalligan | Weir | No | No | Yes | Yes |
| Dinin | Bridge Apron | Yes | No | Yes | Yes |

**Figure 1.**
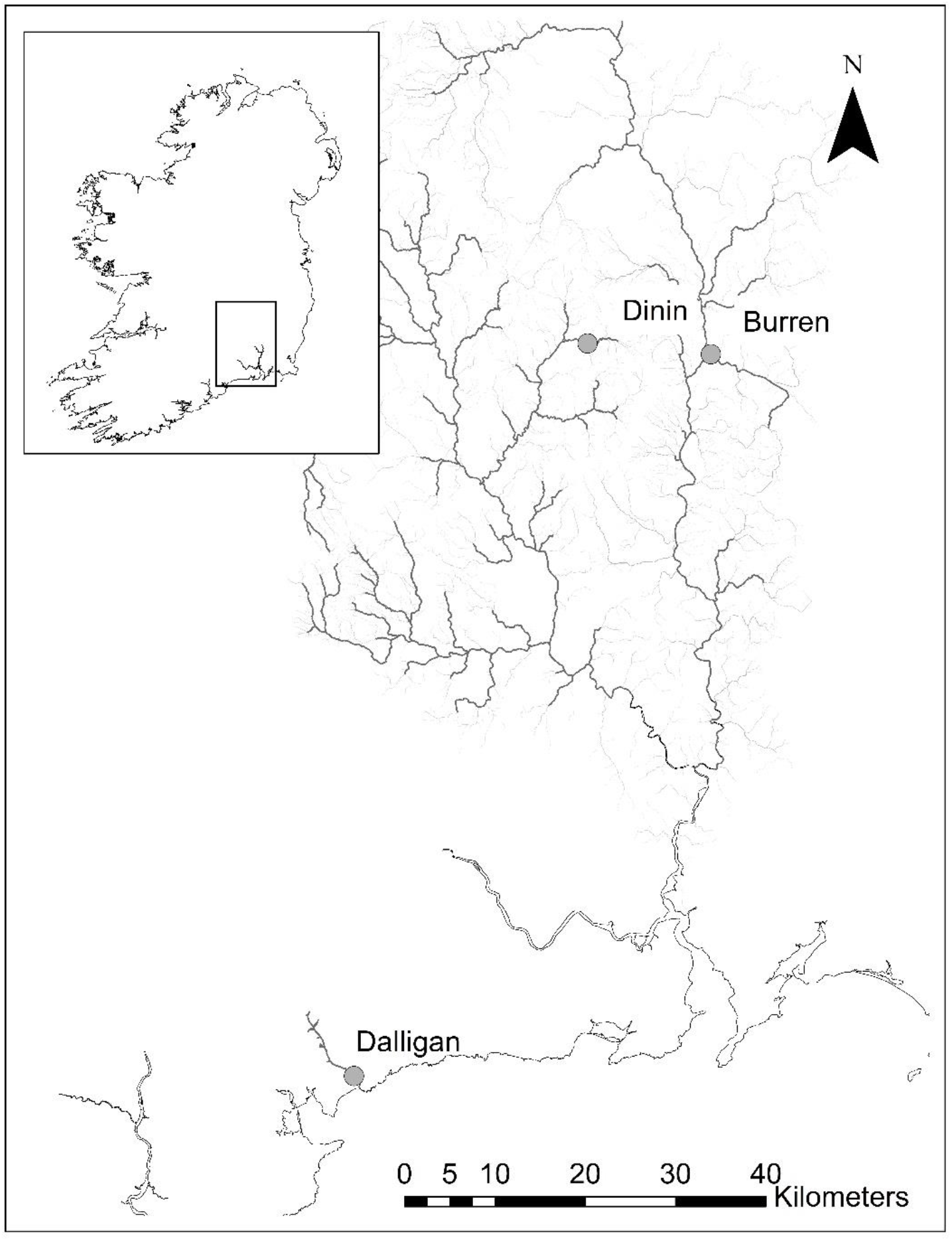
Map showing the locations of the sampling sites in this study.

### 2.3. eDNA collection, filtering and extraction

Environmental DNA samples were collected from each river in sterilized 2L containers, and filtered in the field using a peristaltic pump. Three replicate eDNA samples were collected both upstream and downstream of each river obstacle. One negative field control per location (upstream and downstream) consisting of ddH20 was also filtered, resulting in a total number of six eDNA samples and two field controls collected per river. Environmental DNA was collected on 47 mm glass microfiber filters (1.5 μm) and placed into 2.0 mL Eppendorf tubes prior to being frozen at −20° C. All work with eDNA was carried out in a dedicated Low Copy DNA laboratory to reduce contamination risk. Environmental DNA was extracted using a modified version of the CTAB (cetyltrimethylammonium bromide) protocol (Möller, Bahnweg, Sandermann, & Geiger, 1992). One-half of a glass microfiber filter was placed into a new 2.0 mL Eppendorf tube, to which 750 μL of CTAB buffer (100 mM Tris-HCL, 20 mM EDTA, 1.4 M NaCl, 0.2%, 2% CTAB), and 7 μL of Proteinase K (20 mg mL^−1^) was added. Samples were vortexed for 10 seconds and incubated at 56° C for 2 hours, after which 750 μL of Phenol/Chloroform/Isoamyl Alcohol (25:25:1 v/v) was added. Samples were manually mixed for 15 seconds and centrifuged (11,000 × g, 20 min). The aqueous phase was transferred to a new tube containing 750 μL of Chloroform/Isoamyl Alcohol (24:1 v/v), the manual mixing and centrifugation steps were repeated, and the aqueous phase was transferred to a new tube. The eDNA was then precipitated by adding one volume of isopropanol alcohol to the aqueous phase and incubating the mixture at −20° C for 1 hour, and then centrifuged (11,000 × g, 20 min). The pellets were washed with 750 μL of 70% ethanol and centrifuged (11,000 × g, 5 min). The ethanol was carefully removed, and the pellets dried in a heating block (50° C, 5 min) before resuspending the eDNA in molecular-grade water.

### 2.4. eDNA assay deployment

Environmental DNA concentrations were determined by qPCR using an Applied Biosystems ViiA™ 7 (Life Technologies, Inc., Applied Biosystems, Foster City, CA) quantitative thermocycler. The qPCR reaction was conducted in a final reaction volume of 30μL, comprised of 15 μL of TaqMan® Environmental Master Mix 2.0 (Life Technologies, Applied Biosystems, Foster City, CA), 3 μL of each primer (final concentration of 2 μM), probe (final concentration of 2 μM), DNA template (3 μL) and ddH2O. Warm-up conditions of 50°C for 2 min and 95°C for 10 min, followed by 40 cycles between 95°C for 15 s and 60°C for 1 min were used for the qPCR run. DNA extracted from *S. salar* tissue (quantified with NanoDrop®-1000, Thermo Scientific, Wilmington, DE) was used to generate the standard curve using seven 10:1 serial dilutions. Concentrations for the serial dilution ranged from 3ng/μL to 3 × 10^−6^ ng/μL. The eDNA field samples were run on two separate 96-well clear qPCR plates. Each plate had 3 no-template controls (NTCs) to ensure no contamination occurred during the preparation of the qPCR plate. Individual standard curves were generated for each qPCR plate (y = −3.22x + 19.968, efficiency = 100.018% (1) and y = −3.25x + 20.091, efficiency = 103.101% (2)). All standard curve samples, field samples and controls were quantified in triplicate (three technical replicates). A positive detection was defined as being within the range of the standard curve, and when at least 2 out of the 3 technical replicates contained amplifiable DNA with Cq differences not exceeding 0.5. If the difference between 1 out of 3 technical replicates exceeded 0.5Cq, this technical replicate was excluded from the study. However, if the Cq value of 2 out of 3 technical replicates differed by more than 0.5Cq, the entire sample was excluded from further study. As *S. trutta* was present in all rivers, both upstream and downstream of the obstacles (Table 1), this species was used as a positive field control to test for the presence of amplifiable DNA in sites where no *S. salar* was recorded during electrofishing surveys. The *S. trutta* assay from previously published work (Gustavson et al., 2015) was used on eDNA samples from above the bridge apron in the Dinin river, and above and below the weir in the Dalligan river. Three replicates per location with one technical replicate were used for this analysis.

## 3. Results and Discussion

The present assay was successful in detecting *S. salar* DNA *in silico*, *in vitro* and *in situ*. The assay did not amplify the DNA of closely related species (*S. trutta*) or any other species included in the specificity test. The lowest detected eDNA concentration within the range of the standard curve was 0.016 ng L^−1^ at Cq 34.5 (average over 3 technical replicates, standard deviation 0.0015 ng L^−1^). For the purposes of analysis, one technical replicate from the 1:7 serial dilution was disregarded (equation 1), and the entire 1:7 dilution for the standard curve (equation 2) was disregarded because differences in Cq values between either one or more technical replicates in these samples exceeded 0.5. For the remainder of the samples, however, the standard deviation between technical replicate Cq values ranged from 0.011 to 0.303.

The results of the eDNA analysis mirrored what was observed in the electrofishing surveys. At each site where the presence of *S. salar* was confirmed by electrofishing, its presence was confirmed by eDNA analysis (Table 2, Figure 2). At sites where *S. salar* was not detected by electrofishing, a negative result was also obtained in the eDNA samples when assessed with the *S. salar* assay (Table 2, Figure 2). However, detectable eDNA was confirmed at all sites including the sites where no *S. salar* DNA was detected, as amplification occurred when the same samples were run in qPCR with the *S. trutta* assay. No DNA was amplified in any of the NTCs or negative field controls.

**Table 2.**
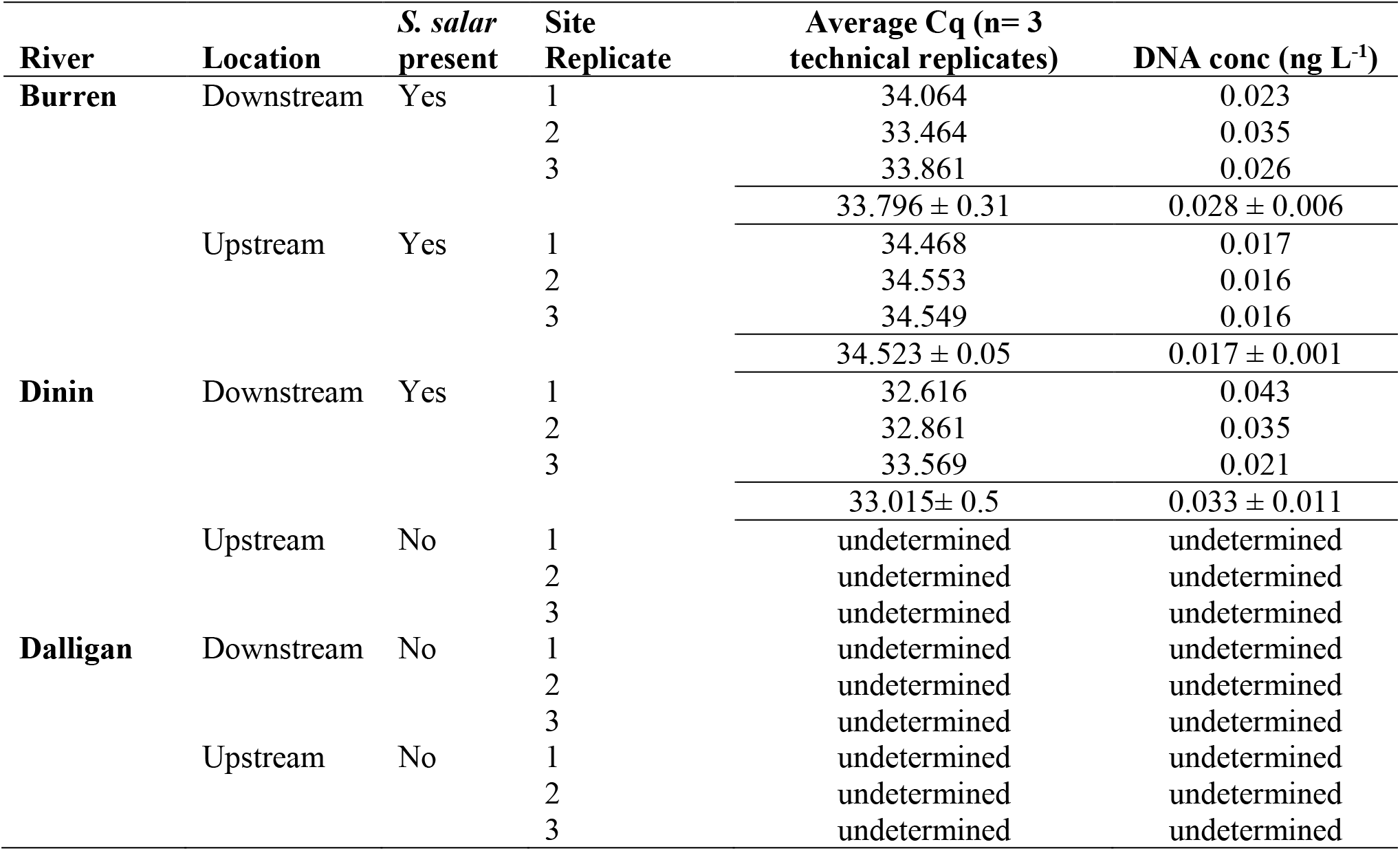
The Cq values and eDNA concentrations (ng L^−1^) (average over three technical replicates per site replicate) from the *S. salar* assay in each river. Average concentrations (± SD) are given for each location (upstream or downstream of the river obstacle).

**Figure 2.**
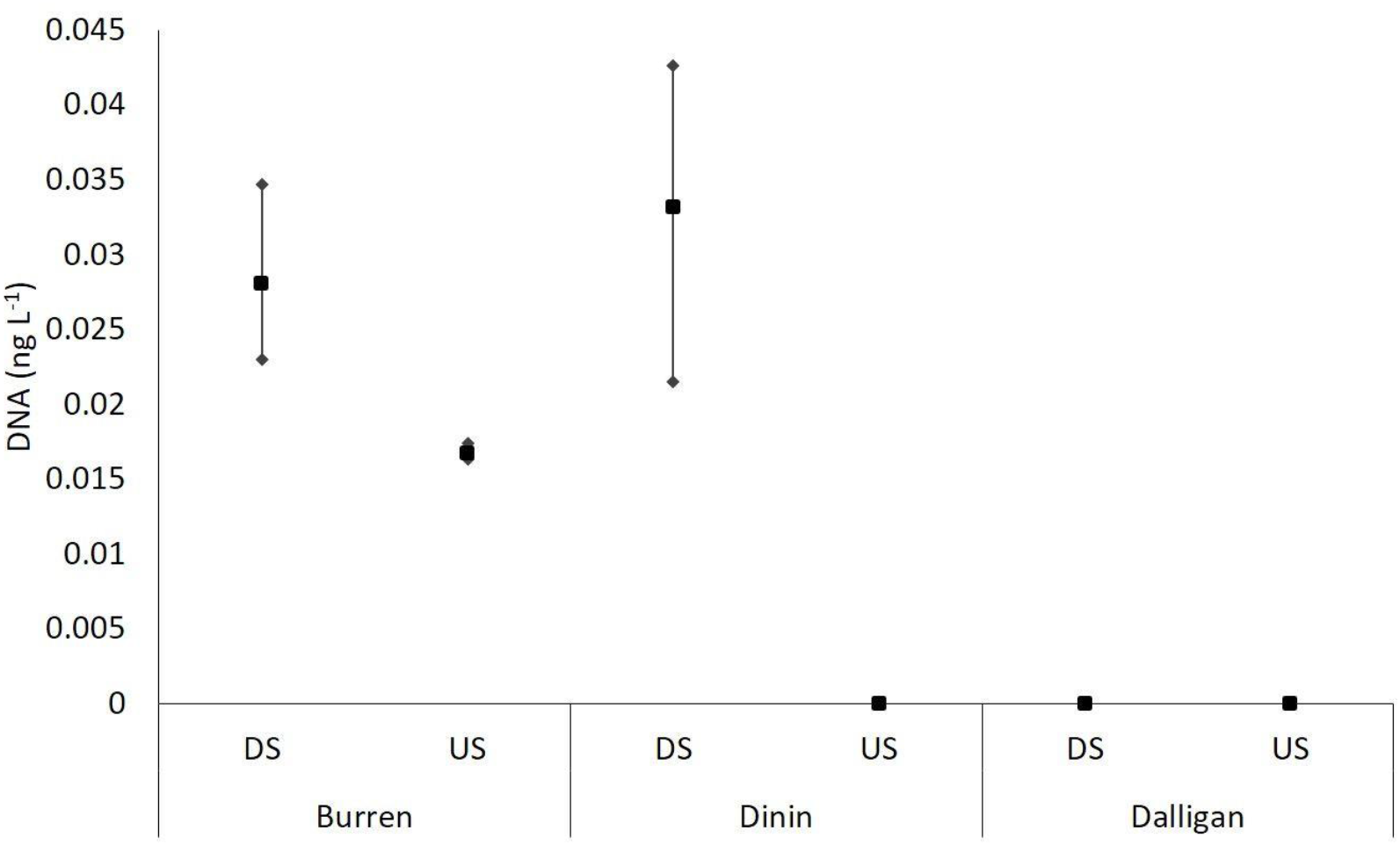
Graph showing the mean and range (maximum and minimum) of *S. salar* eDNA concentrations (ng L^−1^) at each location (downstream (DS) or upstream (US) of the river obstacle) within each river sampled.

The assay successfully confirmed either the presence or absence of *S. salar* in each sampling location, demonstrating the potential future use of this assay for detecting the species without traditional sampling methods. Furthermore, this eDNA assay would be particularly valuable for monitoring *S. salar* year-round. Traditional sampling methods are typically carried out during specific times of the year. For example, redd counts are only possible during the spawning period, and electrofishing surveys are typically restricted to the summer months when water levels are low, and fish are not migrating. While fish counters and traps can provide year-round records of *S. salar* movements, the structures themselves can act as obstacles to the movement of other, non-salmonid fish. For example, resistivity fish counters are typically placed on sloping weir-like structures (Lucas & Baras, 2000) which have been shown to impede the movement of lamprey *Lampetra fluviatilis* L. (Lucas, Bubb, Jang, Ha, & Masters, 2009; Russon, Kemp, & Lucas, 2011) and barbel *Barbus barbus* L. (Lucas & Frear, 1997).

While this study clearly demonstrates the value of eDNA as a tool for monitoring the impact of river obstacles on *S. salar*, it could be applied in numerous different contexts including monitoring *S. salar* escapes from fish farms outside the native range. In addition, the use of eDNA as a monitoring tool could largely reduce the spread of invasive alien species, since eDNA sampling must be carried out with minimum equipment that must be sterilised to avoid DNA contamination (Herder et al., 2014).

To conclude, the assay presented here is an efficient and effective method of detecting *S. salar* in rivers. Similar to Laramie et al. (2015) the assay presented here could be used to identify new conservation areas for the species, and additionally, can provide evidence to support remediation action, for example removing river obstacles that may be preventing the migration of the species.

## Acknowledgements

This project was funded by the Atlantic Salmon Trust (Salmo Slime project) with additional support from the Irish Environmental Protection Agency (Reconnect project). The authors would like to thank Inland Fisheries Ireland for carrying out all the electrofishing surveys which supported this research.

